# Combination of Expert Guidelines-based and Machine Learning-based Approaches Leads to Superior Accuracy of Automated Prediction of Clinical Effect of Copy Number Variations

**DOI:** 10.1101/2022.12.23.521405

**Authors:** Tomáš Sládeček, Michaela Gažiová, Marcel Kucharík, Andrea Zaťková, Zuzana Pös, Ondrej Pös, Werner Krampl, Erika Tomková, Michaela Hýblová, Gabriel Minárik, Ján Radvanszky, Jaroslav Budiš, Tomáš Szemes

## Abstract

Clinical interpretation of copy number variants (CNVs) is a complex process that requires skilled clinical professionals. General recommendations have been recently released to guide the CNV interpretation based on predefined criteria to uniform the decision process. Several semiautomatic computational methods have been proposed to recommend appropriate choices, relieving the clinicians from the tedious search in vast genomic databases. We have developed and evaluated such a tool called MarCNV and tested it on CNV records collected from the ClinVar database. Alternatively, the emerging machine learning-based tools, such as the recently published ISV (Interpretation of Structural Variants), showed promising ways of even fully automated predictions using wider characterization of affected genomic elements. Such tools utilize features that are additional to ACMG criteria, thus, they have the potential to significantly improve and/or provide supportive evidence for accurate CNV classification. Since both approaches contribute to evaluation of CNVs clinical impact, we propose a combined solution in the form of a *decision support tool* based on automated ACMG guidelines (MarCNV) supplemented by a machine learning-based pathogenicity prediction (ISV) for classification of CNVs. We provide evidence that such a combined approach is able to reduce the number of uncertain classifications and reveal potentially incorrect classifications using automated guidelines. CNV interpretation using MarCNV, ISV, and combined approach is available for non-commercial use at https://predict.genovisio.com/.

## Introduction

Broad implementation of array- and massive parallel sequencing (MPS) technologies leads to the unleashing of an ever-increasing genetic variability. This raises the demand for correct understanding and interpretation of the impact of the identified variants on human health, especially in clinical settings. The process of interpretation of genomic variants is complex and in many cases yields ambiguous results, which may even vary between laboratories. In order to alleviate this obstacle, technical standards for the interpretation and reporting of variants have been developed by the American College of Medical Genetics and Genomics (ACMG) and the Association for Molecular Pathology (AMP). The first initiative was developed for small sequence variants ^1^, as they are better understood. These standards are not suitable for the large-scale variants, such as copy number variations (CNVs), though some of the basic rules for the interpretation are consistent across all types of genomic variability. Evaluation of CNVs*’* clinical impact is really challenging since CNVs typically affect multiple functional genomic elements simultaneously and specific rules must therefore be applied.

At present, several approaches are available to clinicians in the decision-making process regarding interpretation of CNVs*’* clinical significance/impact. In the main approach, CNVs can be classified following a recently published joint consensus recommendation of the ACMG and the Clinical Genome Resource (ClinGen) (in brief: ACMG criteria) ^2^, by selecting choices/options and respective score values for individual guideline categories ^3–5^. These professional standards tend to encourage consistency and transparency of CNV evaluation across clinical laboratories. In another approach, CNV classification is based on the fully automated machine learning algorithms that can go beyond the established scheme, thus having a great potential to further improve the prediction accuracy ^6–8^. The CNVs*’* pathogenicity effect can be also evaluated according to aggregation of per-base single nucleotide polymorphism (SNP) pathogenicity scores within the CNV intervals ^9^. One example of a tool that uses this approach is the SVscore ^9^, which aggregates the SNPs*’* pathogenicity scores according to CADD (Combined Annotation Dependent Depletion) ^10^. Recently, another expert system for CNVs*’* clinical impact classification has been presented – ABC system ^11^ consisting of three-step classification: functional grading (A), clinical grading (B), and optional selection of standard variant comment based on a combined class (C). Authors propose to use it independently or as a supplement to the ACMG system.

Approach based on ACMG guidelines follows a scoring scheme divided into five basic sections: 1) Initial Assessment of Genomic Content; 2) Overlap with Established Triplosensitive (TS), Haploinsufficient (HI), or Benign Genes or Genomic Regions; 3) Evaluation of Gene Number; 4) Detailed Evaluation of Genomic Content Using Cases from Published Literature, Public Databases, and/or Internal Lab Data; and 5) Evaluation of Inheritance Patterns/Family History for Patient Being Studied ^2^. In other words, it is based on the genomic content of the identified CNV, namely on the evaluation whether the region that the given CNV covers also features any present protein-coding genes or elements with important functions, known HI/TS genes or known benign CNVs. Detailed evaluations of CNVs*’* genomic content from the published literature, studies, and available databases is also included, together with the family history of the patient. Based on the sub-intervals of the final score, the 5-tier classification system is implemented and the potential clinical impact of the tested CNV is classified as: benign (B) (≤ −0.99), likely benign (LB) (from −0.98 to −0.90), uncertain significance (VUS) (from −0.90 to 0.90), likely pathogenic (LP) (from 0.90 to 0.98) and pathogenic (P) (≥ 0.99).

The main purpose of the guidelines is to assist clinical laboratories in the classification and reporting of CNVs*’* clinical impact, irrespective of the technology used to identify them. However, many CNVs are evaluated as VUS, thus, frequently inconclusive, and the follow-up classification and validation/curation by a clinician is needed (sometimes after several years).

Another inconvenience is the frequent need for a clinical expert to spend a long time browsing various data sources and publications in order to evaluate individual categories. Selection of different options and assignment of a score for each category is not immediate as the subjective judgment of the clinician may influence the score adjustment in several categories. Thus, based on these criteria, different clinicians may arrive at conflicting CNV classifications.

This has driven the effort to create semi-automated tools that can partially eliminate the tedious process of annotating and assigning a certain score for CNVs by automating the search for information related to the genomic content of CNVs from various sources/databases. Examples of such classification tools that help to automate CNV annotation and classification of individual sections of ACMG criteria include AnnotSV ^3^, ClassifyCNV ^4^ and AutoCNV ^5^. These tools slightly differ in the data sources they use, in the categories from five sections of ACMG criteria that they are able to evaluate (Table 1), and in the way they interpret the details of the ACMG criteria. Thus, the subsequent final score of CNV and the classification of its clinical impact by different tools may differ. In order to finalize evaluation, sometimes it is still necessary to search for genomic content of the CNV in the published literature and to manually evaluate the patient*’*s family history.

**Table 1.**
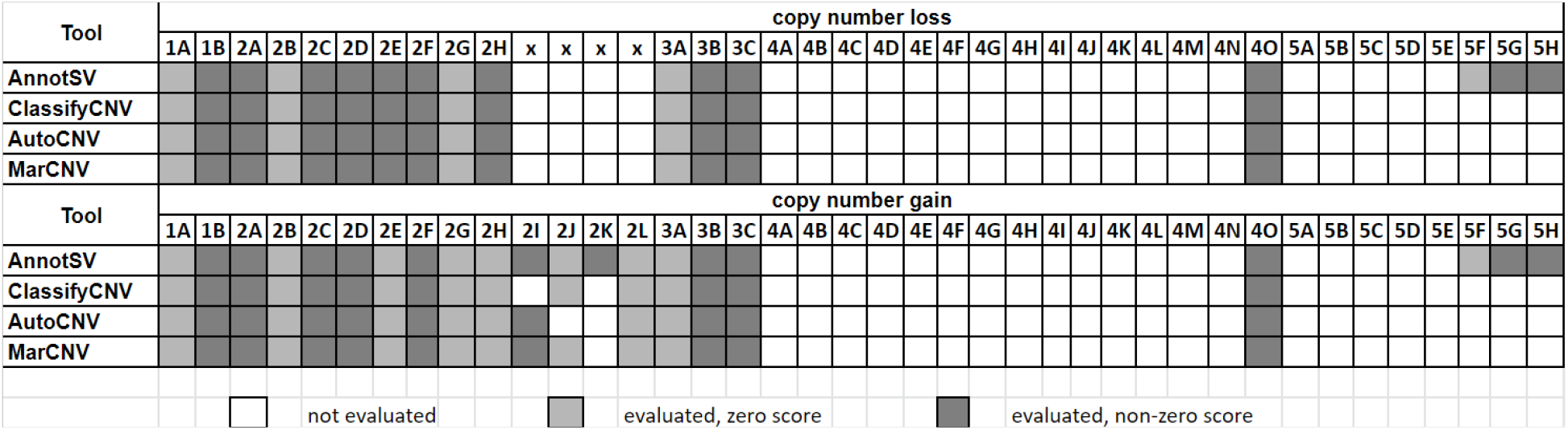
Sections/parameters of the ACMG criteria that are evaluated/not evaluated by the listed semi-automated tools for determining CNVs*’* pathogenicity. The MarCNV tool was designed by us (see methods).

On the contrary, approaches for determining CNVs*’* clinical significance that predict the functional impact of CNVs based on specific trained machine learning algorithms can be fully automated. Several *in silico* CNV pathogenicity prediction tools have been reported recently differing in the specific machine learning algorithms they implement, in the usage of specific genomic elements as features for model training, and in the parameters applied for model training. One of the tools is the ISV (Interpretation of Structural Variants) ^8^ which predicts the probability of pathogenicity of a CNV and its likely clinical impact based on specified CNV coordinates (see the ISV subsection in Methods for details). One of the tools is StrVCTVRE ^6^ focusing on SVs*’* overlapping exons. The model was trained on CNVs collected from the ClinVar database using 17 annotations characterizing gene importance, coding region, conservation, expression, and exon structure as features and is implemented as a random forest. Another available method for pathogenicity prediction is X-CNV ^7^, the model trained using the XGBoost classifier. Recently, we have developed a tool called ISV (Interpretation of Structural Variants) ^8^ which predicts the probability of pathogenicity of a CNV and its likely clinical impact based on specified CNV coordinates (see the ISV subsection in Methods for details).

In this paper, we present an automated tool called MarCNV for evaluation of mainly the first three sections of the ACMG criteria. We provide evidence that both the database choice and the parameters*’* selection influence the resulting score when implemented to some sections of the ACMG criteria. Finally, we evaluated the clinical impact of the selected CNV test sets (from ClinVar database) using MarCNV and ISV (machine learning approach), both individually or together as their combination. We show that the combined approach is able to reduce the number of uncertain classifications (VUS) and reveal the potentially incorrect classifications according to automated guidelines.

## Results

We present MarCNV, an automated tool for evaluation of the first three sections of the ACMG criteria ^2^. We tested the efficacy of MarCNV using several CNV test sets from ClinVar and we then used MarCNV to evaluate the influence of a choice of database parameters on CNVs*’* clinical evaluation and found a way to improve the ACMG criteria. We have managed to join two approaches, namely classification according to the ACMG criteria using MarCNV and automated prediction by ISV, and merge them to create a combined approach (MarCNV + ISV). Moreover, we have demonstrated that this approach not only increases the accuracy of the prediction but also reduces the count of evaluation of CNVs as VUS. We have also performed several comparisons between the individual and combined approaches and compared them with the available ClassifyCNV tool. In the end, using real CNVs from the clinical laboratory practice, we verified and then compared evaluations performed using different approaches.

### Automated ACMG evaluation - MarCNV

#### Comparison with ClassifyCNV

MarCNV is an automated tool for evaluation of mainly the first three sections of the ACMG criteria ^2^. The main difference between MarCNV and ClassifyCNV is the use of different databases for comparison of tested CNVs (Supplementary Table S1). The ACMG criteria are rather strict in evaluation, however, there are still minor differences between MarCNV and ClassifyCNV in their interpretation. For example, to evaluate the overlap with haploinsufficient genes (sections 2C-2E for loss), the ACMG guidelines do not exactly specify which transcript of the gene should be taken into account. MarCNV evaluates all transcripts in the database (and subsequently reports the most severe score), while ClassifyCNV evaluates only the most established transcript. MarCNV cannot distinguish between options 2J and 2K for copy number loss due to unknown patient phenotype, which is a necessary piece of data enabling the decision between these two options. However, it can evaluate option 4O (section 4, *overlap with common population variation*), both for copy number loss and gains. The *accuracy* and *inclusion* of MarCNV and ClassifyCNV are quite comparable (see Fig. 3) while ClassifyCNV evaluated a little bit less CNVs as VUS when compared to MarCNV.

#### The choice of a database of benign CNVs greatly influences the precision of evaluation

We used MarCNV to evaluate some parameters and their effect on the result of the CNV evaluation process. We observed that some sections of ACMG evaluation are very sensitive to the choice of database and its cleaning procedures. As an example, we evaluated the option 2F for CNV losses (2F. *Completely contained within an established benign CNV region*) using different benign CNV databases and filtration settings (Fig. 1). We observed that the different filtration settings worked as a trade-off parameter between the *accuracy* of the prediction and the proportion of evaluated CNVs (*included*). Both parameters varied greatly for tested databases: *accuracy* from 95.92% (GnomAD, no filtration) to 99.02% (DGV-GOLD-INNER, frequency ≥1%); *inclusion* from 15.71% (GnomAD, frequency ≥10%) to 70.96% (DVG, no filtration). As the best option, we have chosen DGV-GOLD-OUTER with the frequency of ≥0.5%, as it has almost the best *accuracy*, while *including* almost a quarter of all CNVs (*accuracy*=98.26%, *inclusion*=27.28%) (Fig. 1).

**Figure 1:**
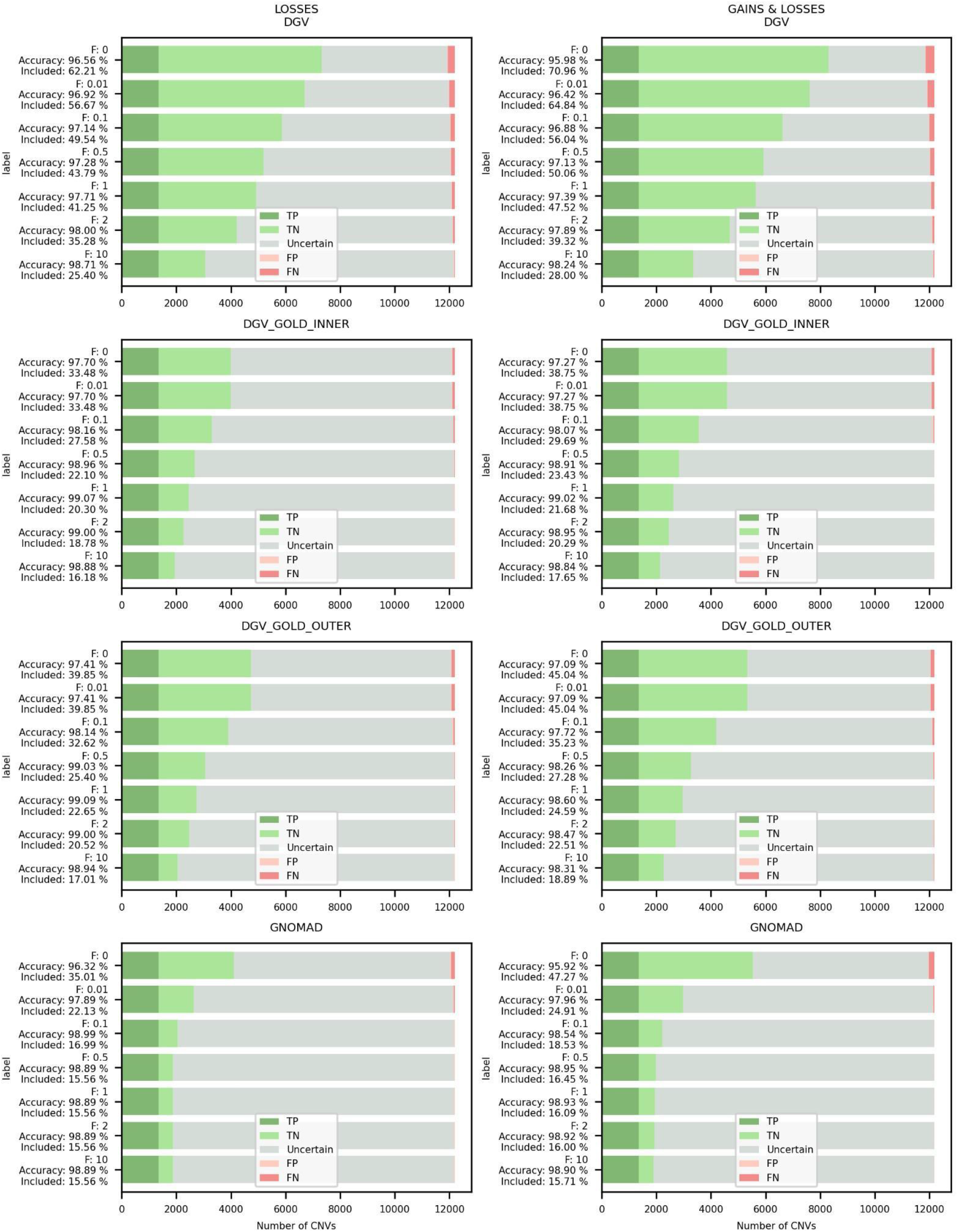
Evaluation of the option 2F for CNV losses by MarCNV using different databases of benign CNVs and their settings. Left column represents analyses performed selecting only “losses” in the search for the *established benign CNV region*, while on the right both “gains & losses*’’* were selected (as recommended by ACMG). GnomAD ^12^ and DGV ^13^ databases that collect common CNVs from large sequencing studies were searched and the results were compared. DGV database was represented in three versions, namely “DGV” that represents all CNVs in DGV database; “DGV-GOLD” stands for “Gold Standard” and represents curated CNVs with INNER or OUTER CNV coordinates. Database search was performed at different minimum population frequency thresholds (“F”) set for a known/established benign variant, with the assumption that a more common CNV is less harmful. The *accuracy* indicates the proportion of correctly evaluated CNVs, whereas *included* represents the percentage of predicted CNVs falling either into the B or P category (see Methods for details). We found the DGV-GOLD-OUTER at population frequency 0.5% to be the best. TP=true positive, TN=true negative, FP=false positive, FN=false negative.

#### Improvement of the ACMG criteria

Using MarCNV, the current ACMG criteria can be adjusted to test for more *accuracy* and/or *inclusion*. For the option 2F for CNV loss we demonstrated that including only benign losses in the database search does not provide any advantage compared to the option chosen by ACMG, where both losses and gains are included. Therefore, in all the following tests we continued with the original ACMG option (Fig. 1).

For options 2A and 2B we demonstrate that inclusion of HI/TS genes with the score 2 or even 1 in the database search (in addition to those with score 3 from the ACMG settings) increased the number of predicted CNVs, while only slightly decreased the accuracy (Fig. 2).

**Figure 2:**
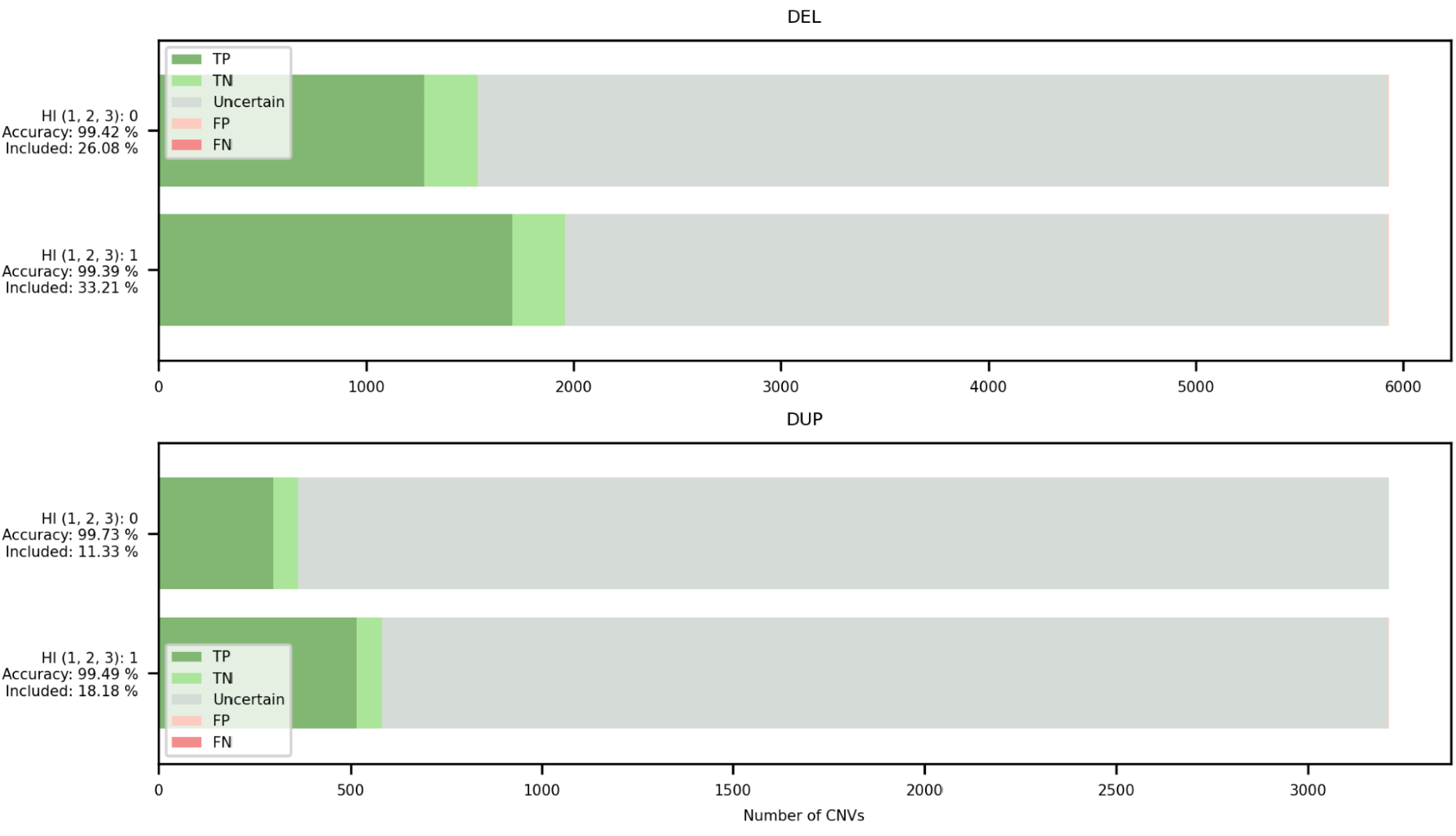
Comparison of options for HI/TS genes for evaluation of the ACMG criteria (sections 2.A and 2.B) - both losses (DEL, upper panel) and gains (DUP, lower panel) were analyzed. “HI(1,2,3): 0” represents an option where only HI/TS genes/regions with score of 3 taken into account (as in regular ACMG guidelines); “HI(1,2,3): 1” represents a search where also HI/TS genes with scores of 1 or 2 were included (together with those with score of 3). The *accuracy* indicates the proportion of correctly evaluated CNVs, whereas *included* represents the percentage of predicted CNVs falling either into the B or P category (see Methods for details). The DGV-GS-OUTER database with frequency ≥0.5% was used for parsing benign CNVs. The x-axis represents the number of CNVs. TP=true positive, TN=true negative, FP=false positive, FN=false negative.

### A novel approach combining MarCNV and ISV

#### Application of the new combined approach can increase *accuracy*

We confirmed that the ACMG scoring scheme is rather strict and the majority of CNVs are classified as VUS. On the other hand, the machine-learning tool ISV is able to classify many more CNVs, however, it does it at the cost of producing more false predictions. We show in Fig. 3 that combined use of these methods yields a superior predictor, with a higher *accuracy* than the raw ISV model, and more predicted CNVs (*included*) than the ACMG scheme alone (see also Supplementary Figure S1 for direct comparison of ClinVar classification of CNVs from the test set and their interpretation using our combined approach).

**Figure 3:**
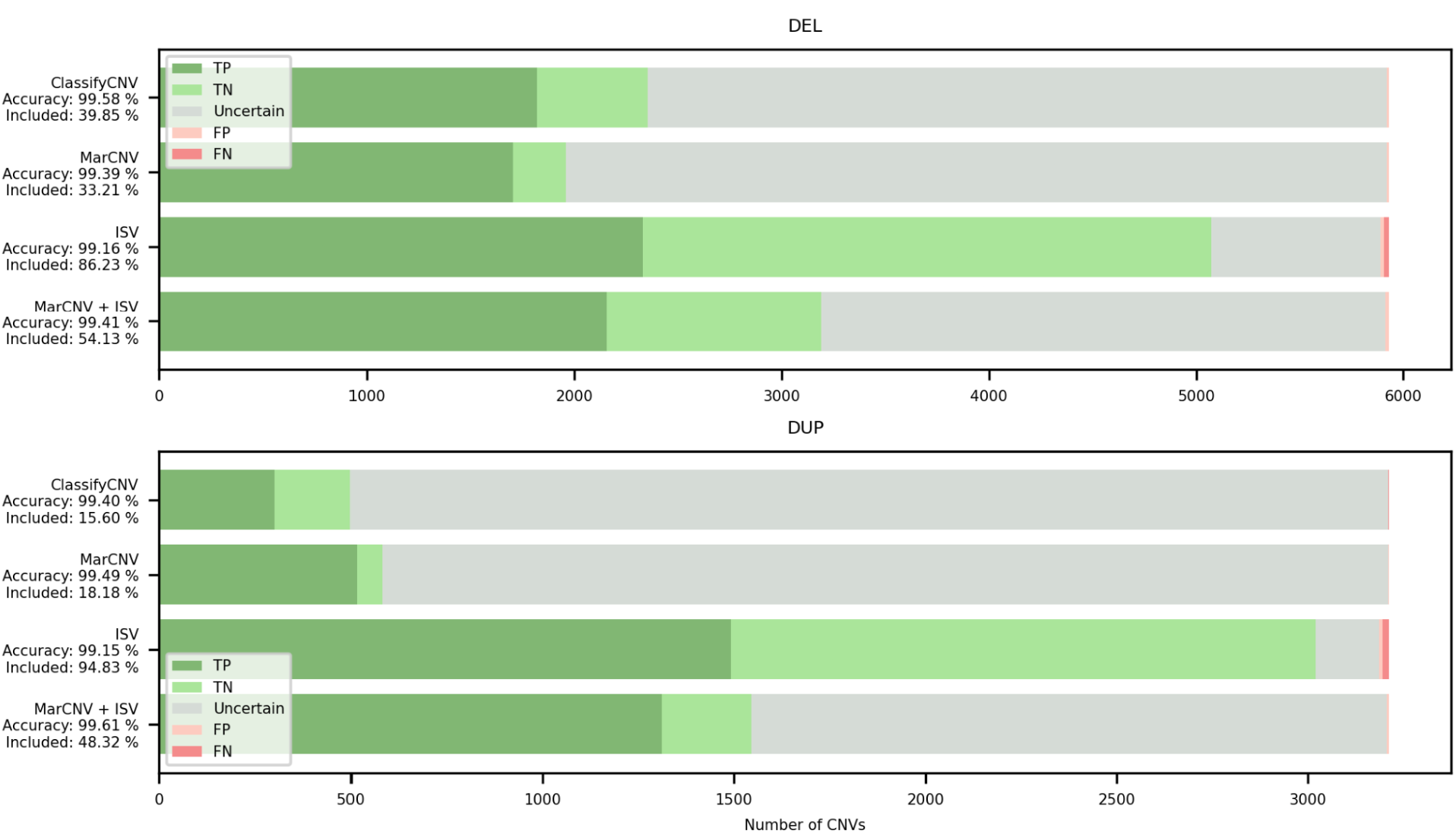
Comparison of ClassifyCNV, MarCNV, ISV and MarCNV+ISV methods of CNV evaluation – both losses (DEL, upper panel) and gains (DUP, lower panel) were analyzed. The *accuracy* indicates the proportion of correctly evaluated CNVs, whereas *included* represents the percentage of predicted CNVs falling either into the B or P category (see Methods for details). The DGV-GS-OUTER database with the frequency of ≥0.5% was used for parsing benign CNVs. The x-axis represents the number of CNVs. TP=true positive, TN=true negative, FP=false positive, FN=false negative.

We observed that *accuracy* of combined approach depends on the MarCNV score and on the ISV ratio (*r*) setting (see Methods), where raising *r* will increase the influence the ISV has on the final prediction. As can be seen in Fig. 4, two ISV ratios are particularly interesting: the first *r ≃* 0.19 and the second *r ≃* 1.99. In both these ratios, we observe a sudden drop in the count of uninformative predictions (gray area). The first drop represents CNVs that had an ACMG score of 0.9, thus, they would normally be classified as LP according to the ACMG scheme. When ISV yields probabilities that are very close to 1, these CNVs will suddenly be classified as pathogenic. This sudden decrease in both ratios is more prevalent for CNV-gains.

**Figure 4:**
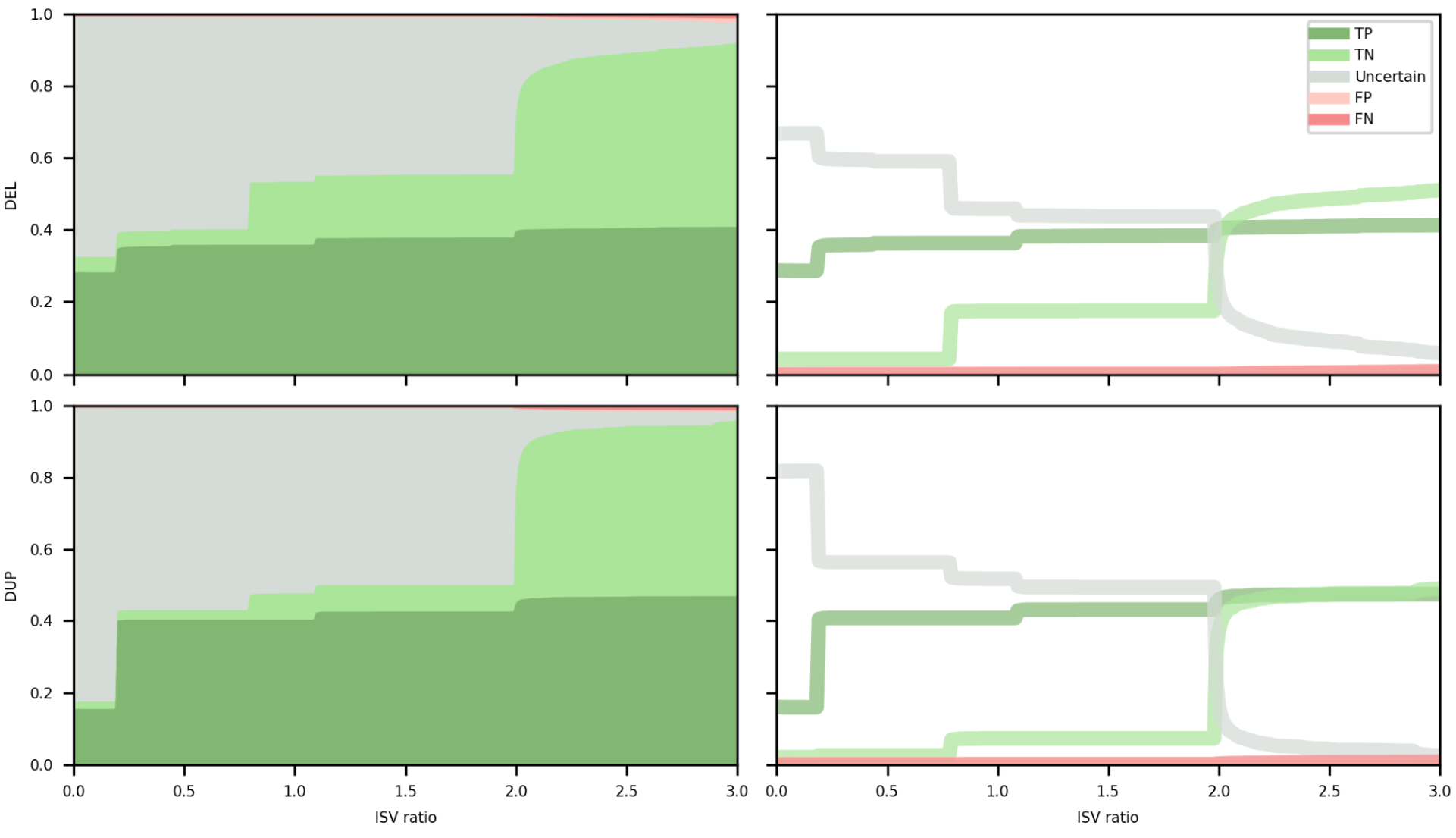
ISV ratio in the combined approach for evaluation of CNV losses (DEL, upper panel) and gains (DUP, lower panel). The left column represents relative counts of True, Uncertain and False predictions, while the right one contains the same information plotted as a line chart individually for each category in order to facilitate comparison of the behavior of individual categories with increasing ISV ratio. A sudden decrease in the count of *uncertain predictions* (gray area) is visible, especially for ISV ratios at *r ≃* 0.19 and *r ≃* 1.99. The Y-axis represents the area (left) or proportion (right) of each individual category at a given ISV ratio (x-axis). Color code for individual categories is the same for both columns and is indicated in the figure (TP=true positive, TN=true negative, FP=false positive, FN=false negative).

For example: If a CNV has ACMG score (MarCNV) of 0.9 and ISV probability of 0.99, then according to the model above with *r* = 0.19, we would get the final score of: 0.9 + 0.19 * (0.99 − 0.5) = 0.9931, which can be classified as P according to the ACMG guidelines. This is partly caused by the overconfidence ISV has in its predictions. Most CNVs usually have either very high or very low ISV probabilities of pathogenicity and that is why the ratio *r* = 0.19 causes the sudden jump.

The second jump comes at ratio *r* = 1.99, where most CNVs with ACMG score close to 0 (VUS), became classifiable after addition of the ISV term. Again, this stems from the fact that if ISV thinks that a CNV is B, it will be very confident and return very low probabilities (around 50% of the predicted B CNVs have ISV probabilities lower than 0.02).

Similar analysis for the ratio 0.19 and 1.99 was performed on clinical samples which will be described below (see also Supplementary Figure S2), where the reduction of CNVs classified as VUS was also confirmed (for r = 1.99).

#### The combined approach leads to an increased number of informative CNVs (falling either into the B or P category *included*) compared to ACMG

Compared to the classification following the ACMG criteria, using a combined approach, we can increase the parameter *included* the number of informative CNVs (falling either into the B or P category). Fig. 4 shows also a reduction of CNVs evaluated as *uncertain predictions* (other than B or P, gray area) when increasing the ISV ratio. When the ACMG criteria (MarCNV) with minimal ISV ratio are used, there are quite a lot of CNVs evaluated as *uncertain predictions*. With increasing ISV ratio in the combined approach, decrease in the number of CNVs evaluated as *uncertain predictions* can be observed. However, with the decreasing CNV*’*s evaluation as *uncertain*, there is an increase in false predictions.

#### Verification of the combined approach on the real samples from the clinical laboratory

We used our combined approach (MarCNV + ISV) to reanalyze 63 monoallelic CNVs evaluated previously by the clinical laboratory (clinical interpretation = CI). CNV classification was performed using standard clinical approach, based on searching in publicly available databases, or in-house/internal databases, as well as the frequency of CNV in publicly available databases. Patient*’*s phenotype was also included in the evaluation. CNVs*’* coordinates and evaluations of clinical significance can be found in Supplementary Table S2.

Nine P CNVs, one B CNV and 37 VUS were assigned to the same category by clinicians as well as by our combined approach. Two LP CNVs, as classified by CI, were evaluated as P using the combined approach. Similarly, seven LP and three LB were classified as VUS, and one VUS as P (see Supplementary Table S2). Moreover, none of the CNVs was evaluated incorrectly – LP/P to LB/B or vice versa. Fig. 5A shows a comparison of the classifications of these CNVs using CI, using MarCNV only and using the combination of MarCNV + ISV (combined approach). We observed that evaluation according to the ACMG criteria (using MarCNV) is quite conservative (compared to CI), since most of CNVs classified as LP and LB using CI were classified as VUS by MarCNV. Adding the ISV evaluation (combined approach) resulted in a reduction of conservative CNV classification by MarCNV: three LP CNVs were changed to P and one VUS CNV was changed to LP.

**Figure 5 A, B:**
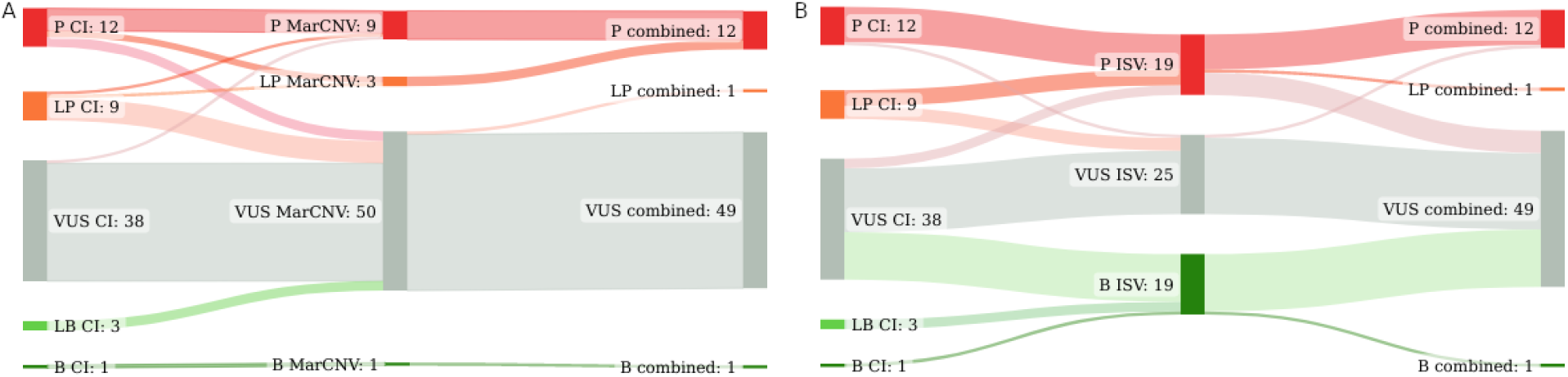
Change in the number of CNVs (obtained from the clinical laboratory) assigned to individual categories using different approaches. The middle column displays the evaluation performed by MarCNV only **(panel A)** (classification according to the ACMG criteria); or by ISV only **(panel B)** (machine learning approach). In both panels, the first column represents evaluation by clinicians (CI = Clinical Interpretation) and the third one evaluation by the combined approach (MarCNV+ISV). Line thickness also illustrates the number of CNVs in individual categories (listed in the Supplementary Table S2. B=benign, LB=likely benign, VUS=variant of uncertain significance, LP=likely pathogenic, and P=pathogenic.

Fig. 5B shows a similar comparison between CI of CNVs, and classification performed by ISV only and by combination of ISV+MarCNV (combined approach). There was a match between CI and ISV evaluation for 11 P, 20 VUS and one B CNVs. One P CNV (as by CI) was evaluated as VUS by the ISV tool, and five LP CNVs were evaluated as P by the ISV tool. After adding MarCNV to ISV evaluation (combined approach), the evaluation became more conservative, since 49 CNVs were evaluated as VUS, while ISV alone predicted only 25 CNVs as VUS.

Direct comparison of the CI was also performed with combined approach, setting the ISV ratios to 0.19 and 0.99 representing two interesting thresholds, at which a significant reduction in CNVs*’* classification as VUS occurs (see also section *‘*Use of a new combined approach can increase accuracy*’* for more details about these ISV ratios) (Supplementary Figure S2; Supplementary Table S2).

Taken together, our results indicate that both the CI and evaluation according to the ACMG criteria (using MarCNV) lead to the assignment of many CNVs to the VUS category. In comparison, the ISV prediction tool changed the predictions of 18 CNVs that clinicians classified as VUS into B (15 CNVs) or P (3 CNVs). However, this may produce some false predictions.

When we compared the results of CNVs*’* clinical significance evaluation between MarCNV and ISV (Supplementary Figure S3), we observed that MarCNV evaluates more CNVs as VUS in comparison to ISV, and no CNV is classified as LB or LP by ISV.

#### Conflicting Predictions

Six non-overlapping CNVs were found in the testing datasets, for which both MarCNV and ISV produced predictions contradicting the ClinVar labels (losses of chr22:18750783-25518625, chrX:155239112-155691896, chrX:77775163-77949550, chr1:16951500-17071209, and chr1:145138148-146401981, and gain of chr2:7495123-87705899). The above-mentioned CNVs either overlap with an established haploinsufficient gene/genomic region, and/or contain known pathogenic CNVs. Therefore, MarCNV and ISV classified them as P, regardless of the B ClinVar label. Moreover, some of these CNVs have become obsolete and are no longer valid in the dbVar database. Conflicting predictions of individual CNVs are described in more detail in Supplementary Notes 1.

## Discussion

Expert-designed ACMG criteria for CNV classification represent a comprehensive scheme and are already widely used by clinicians. These ACMG and ClinGen standards for interpretation and reporting of CNVs provide a semi-quantitative point-based scoring system based on the most relevant categories of evidence, such as included genomic content, dosage sensitivity predictions and curations, predicted functional effect, clinical overlap with patients in the medical literature, evidence from case and control databases, and inheritance patterns of individual CNVs ^2^. Using the scoring metrics, a laboratory geneticist should assign any CNV reported in a patient to one of five main classification categories ^2^.

Most sections of the ACMG criteria can be analyzed by automated processes and several tools were created for this purpose, including MarCNV presented in this paper. Such computational methods recommend appropriate selection of criteria, relieving the clinicians from tedious search in vast genomic databases. We evaluated MarCNV and compared it to previously reported ClassifyCNV using several test sets of CNV records collected from the ClinVar database. This approach enabled us to confirm their comparable performance.

The ACMG criteria assess the pathogenicity of a patient*’*s CNV taking into account also their phenotype, inheritance pattern and family history, which is not always possible, mainly due to the unknown phenotype and family history. MarCNV is able to evaluate mainly the sections dealing with the genomic location of the evaluated CNV, not inevitably the clinical manifestation in the patient, thus, it works mainly for sections 1-3 of the ACMG scheme.

We used the MarCNV tool to evaluate some parameters and their effect on the result of the CNV evaluation process. We observed that some sections of the ACMG evaluation are very sensitive to the choice of database and its cleaning procedures. As an example of this process, we changed the database of *known benign CNVs* for evaluation of sections 2F-G for loss and 2C-2G for gain and evaluated the ACMG criteria. The best performance was obtained using the DGV database curated with frequency ≥ 0.5% and outer coordinate representation, as it has almost the best *accuracy*, while including almost a quarter of all CNVs (*accuracy*=98.26%, *inclusion*=27.28%). The choice of the database is important for *accuracy* and *inclusion*, since if the CNV is recognized as 2F and 4O (for loss) based on the selected database of benign CNVs, it can already be interpreted as a score that is negative enough to evaluate such CNV as benign.

We also show that the current ACMG criteria can be adjusted to allow for more *accuracy* and/or *inclusion*, as well as improve the CNV evaluation. We found out that inclusion of HI/TS genes with lower scores increased the number of predicted CNVs, while it only slightly decreased the *accuracy*. We continue testing additional improvements to the ACMG criteria and implement them to MarCNV in a way enabling the evaluation of any given CNV with “standard” or improved ACMG criteria.

In the process of interpretation of small DNA sequence variants ^1^ several *in silico* tools are widely used. Similar fully automated tools, such as ISV ^6–8^, that use prediction methods based on machine learning algorithms to evaluate the clinical impact of CNVs have also been developed.

Output file of the classification according to the ACMG criteria contains evaluation in five sections showing also which of the criteria was selected as well as points acquired to contribute to the final evaluation of clinical significance. In the machine learning approach, the effect of individual attributes that contributed to the pathogenicity prediction can be viewed using the SHAP values of each attribute.

To classify clinical impact of CNVs, ACMG scoring scheme uses genomic content of the CNV, such as protein-coding, haploinsufficient and benign genes, functionally important elements that are overlapped by the CNV, as well as any overlap with known benign CNVs. The machine learning approach predicts the CNV*’*s clinical significance mainly based on gene annotations, such as protein-coding genes, conservation and their importance, regulatory elements. Moreover, it considers gene expression and exon structure.

The ACMG scoring scheme is rather strict when it comes to classifying CNVs. We observed that the majority of CNVs were classified as VUS. On the other hand, ISV, the machine-learning tool was able to classify many more CNVs, however, at the cost of producing more false predictions.

Evaluation of the advantages of both mentioned approaches lead us to propose here to merge them into a combined method by combining the CNV clinical impact prediction tools (ISV) with the current scoring system of the ACMG and ClinGen standards (MarCNV). In the presented work, we used selected test sets of CNVs from the ClinVar database to evaluate and compare each individual approach (MarCNV and ISV) to a combined method (MarCNV + ISV). We provide evidence that by employing a combined approach, we can increase *accuracy* of predictions as well as the number of informative (falling either into the B or P category) CNVs compared to using the ACMG criteria only. In this way, a more precise evaluation of CNVs*’* clinical impact can be achieved.

*Accuracy* of our combined approach depends on the MarCNV score and the ISV ratio (*r*) setting. In general, the higher the ISV ratio, the greater the influence ISV has on the final prediction. We tested the effect of the increasing ISV ratio only for the range from 0 to 3, since with higher ISV ratios the result will depend mostly on the ISV tool. When *r* = 1, ISV alone will not have the power to decide the CNV outcome where the ACMG score is between -0.48 and 0.48.

Our combined approach was verified also on the real samples from a clinical laboratory. This analysis confirmed that evaluation according to the ACMG criteria (MarCNV) is quite conservative (compared to classification by clinicians), but the combination of both approaches (including ISV) resulted in a reduction in the number of *uncertain predictions* and an increase in the number of CNVs assigned to the category P or B. However, the combined approach may also lead to some false predictions.

It is important to remember that CI includes also patient and family anamnesis, while automated tools such as MarCNV as well as ISV use as an input the CNV*’*s coordinates only. However, after applying the combined approach, if necessary, clinicians can still contribute significantly by evaluation of the patient*’*s anamnesis and including their data.

In conclusion, we can state that the combined application of the ACMG scoring scheme (MarCNV) and a machine learning-based tool (ISV) leads to more robust predictions. We believe that this approach may provide valuable evidence supporting the decision-making process during evaluation and classification of CNVs, especially in clinical settings. The current ACMG standards and guidelines for interpretation of sequence variants (especially of those developed for SNVs) do consider *in silico* predictions as supporting evidence, although with a low weight. However, machine-learning tools for CNV evaluation, such as ISV, were developed almost simultaneously with the proposal of the ACMG standards for CNVs, thus, they could not have been inserted into the ACMG criteria scheme previously. Since there has been a significant improvement in such *in silico* prediction tools recently, we believe, they could be even included as supporting evidence with more power/weight in the future revisions of the ACMG scheme.

## Methods

### MarCNV

MarCNV is a new automated tool for evaluation of mainly the first three sections of the ACMG criteria ^2^ (Table 1). It is similar to the already published ClassifyCNV ^4^ and AutoCNV ^5^, which, however, were not available at the time of the implementation.

### ISV

ISV (Interpretation of Structural Variation) is a machine learning-based approach in which the pathogenicity of a candidate CNV is assessed by observing the counts of overlapped subcategories of genes and regulatory elements ^8^. ISV is based on boosted trees and it classifies most variants reliably and with a high accuracy (∼98%) among the compared tools ^8^.

### CNV data sets

For evaluation and training of hybrid models, we collected CNVs from the ClinVar database ^14^ (downloaded on April 27^th^, 2021). Briefly, we extracted unique copy number loss and copy number gain variants with inner coordinates lifted to GRCh38 using UCSC LiftOver tool ^15^. We filtered out all CNVs shorter than 1KBp and longer than 5 MBp. We included only single-copy events denoted by multiplicity of 1 and 3 for losses and gains, respectively. The selected CNV records were then randomly divided into three disjunct data sets: *Training* – 70% (loss: 8533, gain: 6589), *Validation* – 15% (loss: 1829, gain: 1411) and *Testing basic* – 15% (loss: 1829, gain: 1412).

We also prepared two additional test sets using CNVs with more significant impact: *Testing >5MBp)*, including CNVs longer than 5MBp (loss: 1859, gain: 1328); and *Testing multiple*, including CNVs with different multiplicities (multiplicities 0 for loss and 4 for gain) (loss: 2244, gain: 472). Plots and tables presented in the Results section come from concatenation of the following three datasets: *Testing basic, Testing >5MBp, and Testing multiple*. The referred datasets overlap with those used by ISV ^8^.

### Testing the effect of database selection on the CNV evaluation precision

Automatic evaluation tools search for an overlap of the tested CNV in the selected databases that collect common CNVs from large sequencing studies. We used MarCNV to test whether the selection of different benign CNV databases and the settings used during selection can influence the result of automated CNV evaluation. Namely, we compared evaluation outputs of the sections 2F-G for loss and 2C-2G for gain obtained using two databases of known benign CNVs: GnomAD database ^12^, and Database of Genomic Variants of all benign CNVs (DGV) ^13^. Three versions of DGV databases were compared: “DGV” that represents all CNVs in DGV database; “DGV-GOLD” that stands for “Gold Standard” and represents curated CNVs with INNER or OUTER CNV coordinates. In addition, analyses were performed at different population frequency thresholds, with the assumption that more common CNVs are less harmful. Namely, each of the databases was filtered to screen out the CNVs with less than 0.01%, 0.1%, 0.5%, 1%, 2% or 10% frequency respectively. We thus obtained 28 different databases/settings of benign CNVs for evaluation and comparison. We then calculated *accuracy* of the prediction and the proportion of *included* CNVs, as described in the following paragraph.

### Calculation of the “*accuracy*” and “*included*” parameters used in figures

*Accuracy* was calculated as the number of correctly predicted CNVs (green), divided by the sum of correctly and incorrectly predicted ones (considering ClinVar evaluation as the *ground truth*). The “*Included*” parameter means the percentage of the decided/evaluated CNVs, i.e., those predicted either as B or P. Since classes LP and LB only indicate definitive classification without sufficient conclusive evidence to classify as P or B, they are represented as *uncertain predictions* in our analysis. Only CNVs with score >0.99 and < -0.99 were considered as P and B, respectively.

### Improvement of the ACMG criteria

In order to evaluate the effect of other parameters on the result of the CNV evaluation we tried to tweak the current ACMG criteria by changing some settings of the database search: 1) according to the ACMG criteria, both benign losses and gains are included in the database overlap search for section 2F – we performed an analysis with settings that included only benign losses (which is more intuitive, since CNV losses are evaluated); 2) according to the ACMG criteria, only HI and TS genes with score 3 are included in the search for sections 2A and 2B – we tested how the model performs if HI/TS genes with scores 1 and 2 are included.

### A combined approach that uses MarCNV and ISV

The joint combined MarCNV + ISV approach model can be easily represented by the following equation: *f* _*joint*_ *= ACMG* _*score*_ *+ r * (ISV* _*probability*_ *− 0*.*50)*, where *ACMG*_*score*_ is the score from an ACMG scheme (e.g. MarCNV), and *ISV* _*probability*_ is the probability returned by the ISV model (ranges from zero to one). Subtracting 0.5 from the ISV probability will decrease the final score for benign CNVs and increase the score for predicted pathogenic CNVs by 0.50 at most. Calculating the score in this way preserves the ranges set by ACMG.

Coefficient (ISV ratio) scales the contribution that ISV has on the final prediction. If set to zero, the *r* predictor will be identical to the provided ACMG scheme and raising it will increase the influence of ISV. Effect of the increasing ISV ratio from 0 to 3 on the CNV evaluation in the combined approach was tested. For *r* = 3, the ISV probability (minus 0.50 to achieve negative values for benign predictions) would be scaled by a factor of 3.

## Data availability statement

The raw and processed data required to reproduce the above findings are available to download from https://github.com/tsladecek/cnv_interpretation/tree/master/data

## Code availability

Scripts for reproducing results are available at https://github.com/tsladecek/cnv_interpretation. CNV evaluation using MarCNV only, using ISV only, and using combination of MarCNV + ISV (combined approach) is available at https://predict.genovisio.com/ for non-commercial use.

## Supplementary material

**Supplementary Figure S1**. Comparison of clinical significance of the CNVs test set from the ClinVar database (left) against the evaluation obtained by the combined approach proposed in this paper (MarCNV+ISV) (right). Line thickness illustrates the CNV count. The numbers of CNVs assigned to individual categories are shown. B=benign, LB=likely benign, VUS=variant of uncertain significance, LP=likely pathogenic, and P=pathogenic.

**Supplementary Figure S2:** Change in the number of CNVs (obtained from a clinical laboratory) assigned to individual categories using clinical interpretation (CI) and the combined approach (MarCNV+ISV), at ISV ratios set to 0.19 (panel A) and 1.99 (panel B). As described in the main text (section *‘*Use of a new combined approach can increase accuracy*’*), these are two ratios at which a reduction in the number of CNVs classified as VUS occurs (see also Fig. 4 in the main text). Line thickness illustrates the CNV count. The numbers of CNVs assigned to individual categories are shown. B=benign, LB=likely benign, VUS=variant of uncertain significance, LP=likely pathogenic, and P=pathogenic.

**Supplementary Figure S3**. Comparison of the number of CNVs (obtained from the clinical laboratory) assigned to individual categories by MarCNV and ISV. Line thickness illustrates CNV count. The numbers of CNVs assigned to individual categories are shown. B=benign, LB=likely benign, VUS=variant of uncertain significance, LP=likely pathogenic, and P=pathogenic.

**Supplementary Table S1.**
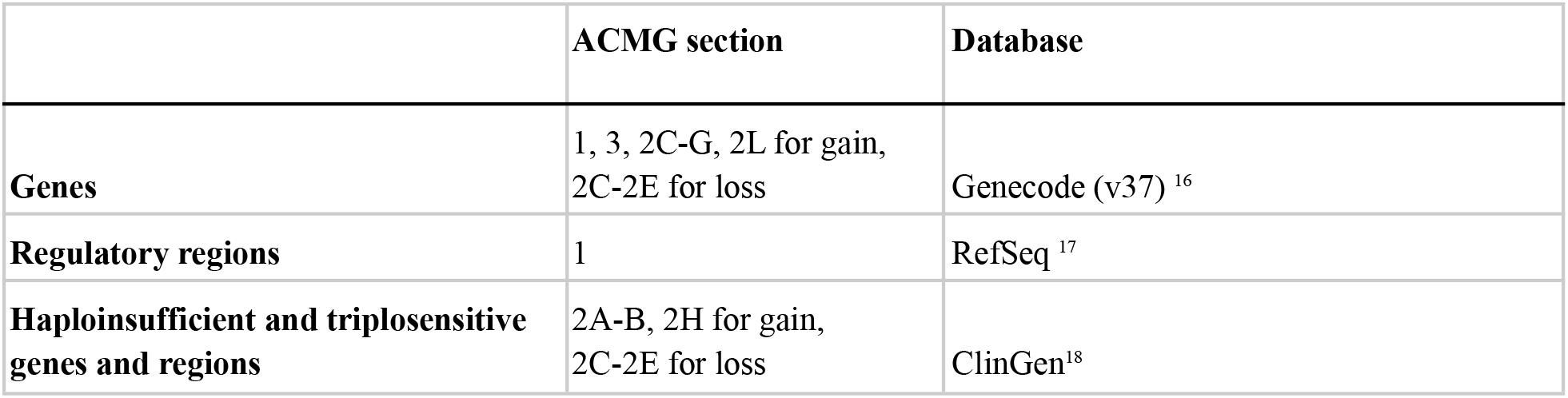
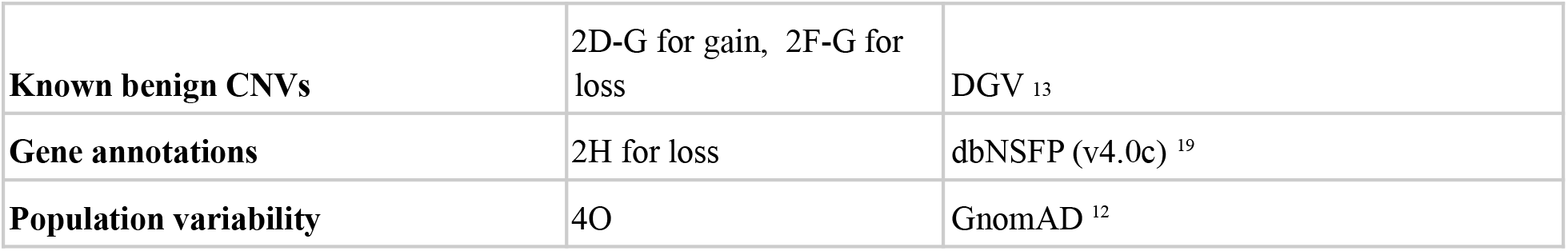
Set of the public databases used by MarCNV in the process of automated CNV evaluation.

**Supplementary Table S2:** Summary of monoallelic CNVs identified in 63 patients from clinical laboratory. CNV type and coordinates are shown, as well as original Clinical Interpretation (CI), together with interpretations performed by MarCNV, ISV and the combined approach. “*r*” represents the ISV ratio set during the evaluation. *r* =1 is the default ISV ratio.

**Supplementary Notes 1: Details on conflicting predictions**

