## Supplementary Figures S1, S2, S3 for "Combination of Expert Guidelines-based and Machine Learning-based Approaches Leads to Superior Accuracy of Automated Prediction of Clinical Effect of Copy Number Variations"

**Supplementary Figure S1.** Comparison of clinical significance of the CNVs test set from the ClinVar database (left) against the evaluation obtained by the combined approach proposed in this paper (MarCNV+ISV) (right). Line thickness illustrates the CNV count. The numbers of CNVs assigned to individual categories are shown. B=benign, LB=likely benign, VUS=variant of uncertain significance, LP=likely pathogenic, and P=pathogenic.

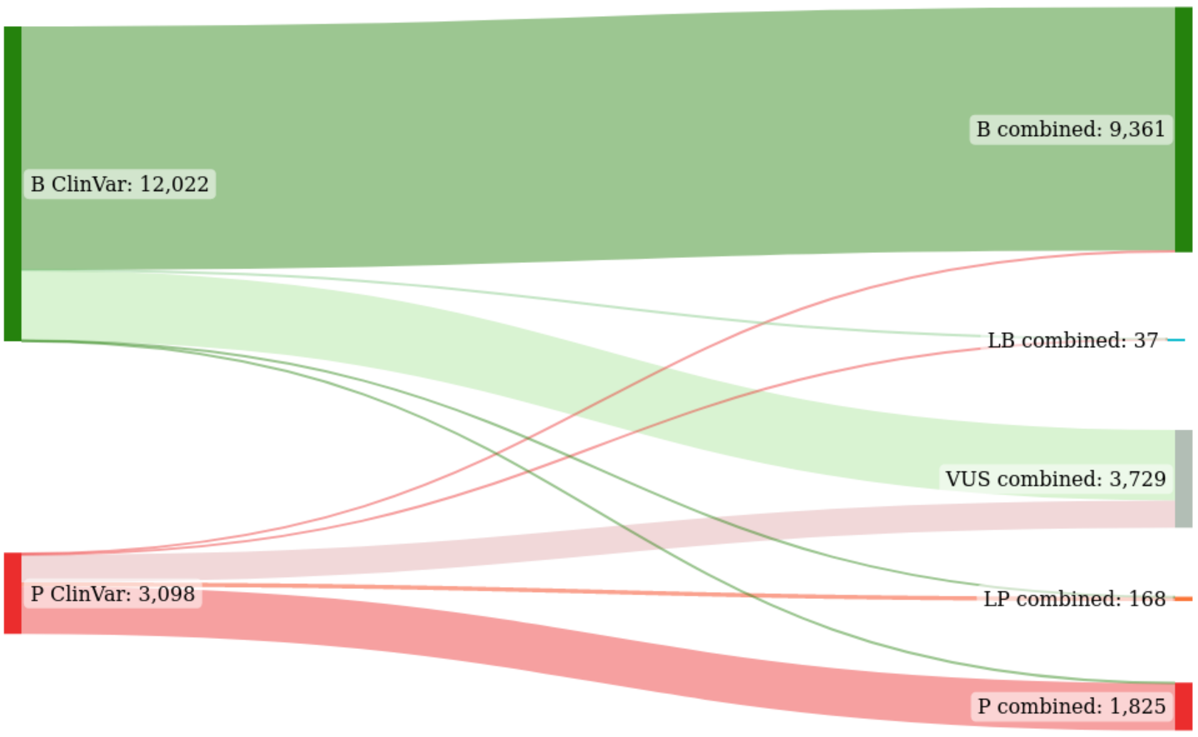


**Supplementary Figure S2:** Change in the number of CNVs (obtained from a clinical laboratory) assigned to individual categories using clinical interpretation (CI) and the combined approach (MarCNV+ISV), at ISV ratios set to 0.19 (panel A ) and 1.99 (panel B). As described in the main text (section 'Use of a new combined approach can increase accuracy'), these are two ratios at which a reduction in the number of CNVs classified as VUS occurs (see also Fig. 4 in the main text). Line thickness illustrates the CNV count. The numbers of CNVs assigned to individual categories are shown. B=benign, LB=likely benign, VUS=variant of uncertain significance, LP=likely pathogenic, and P=pathogenic.

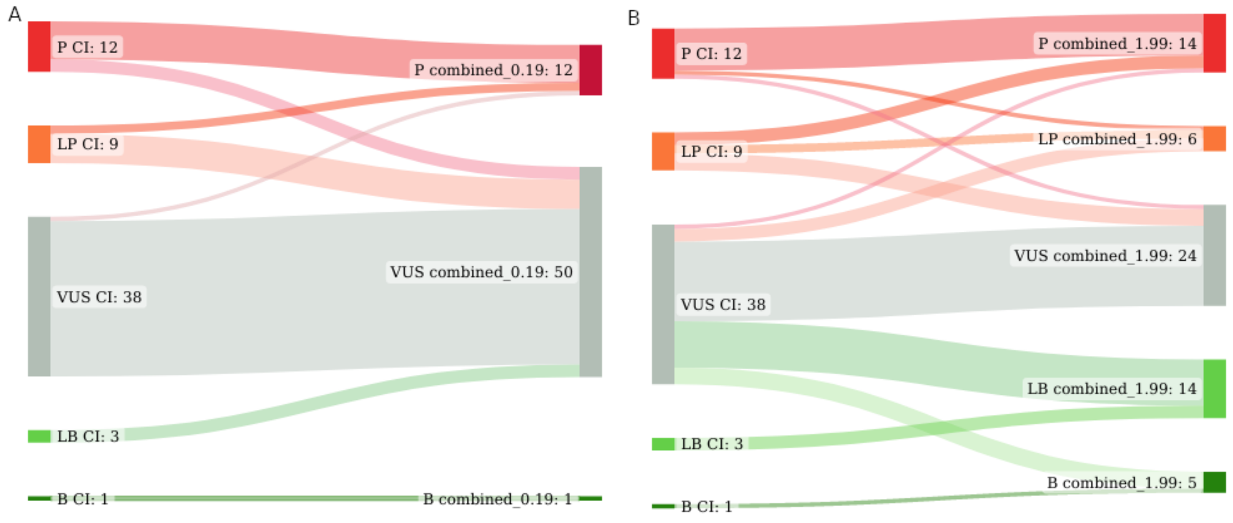


**Supplementary Figure S3.** Comparison of the number of CNVs (obtained from the clinical laboratory) assigned to individual categories by MarCNV and ISV. Line thickness illustrates CNV count. The numbers of CNVs assigned to individual categories are shown. B=benign, LB=likely benign, VUS=variant of uncertain significance, LP=likely pathogenic, and P=pathogenic.

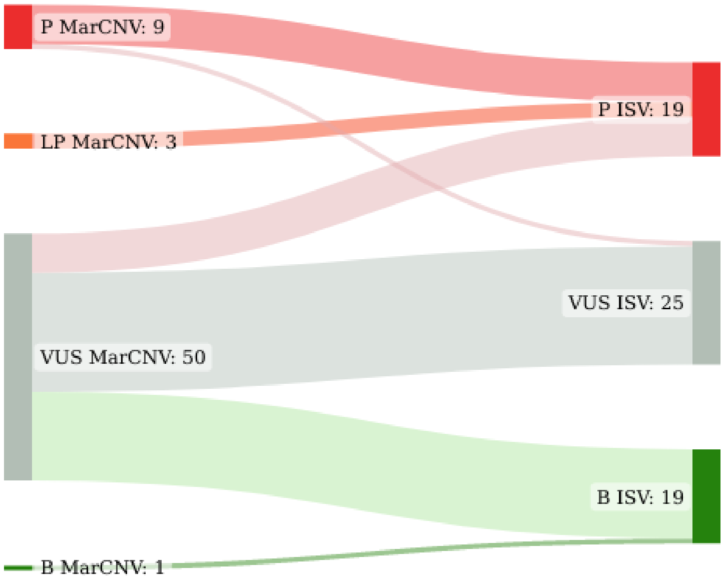
