## Supplementary Notes 1 for "Combination of Expert Guidelines-based and Machine Learning-based Approaches Leads to Superior Accuracy of Automated Prediction of Clinical Effect of Copy Number Variations"

### Supplementary Notes 1 - Details on conflicting predictions

Six non-overlaping CNVs were found in the testing datasets, for which both MarCNV and ISV produced predictions opposite to ClinVar labels:

**Loss of Chr22:18750783-25518625** includes several well-characterized haploinsufficient regions (associated with DiGeorge syndrome #188400). Only one ClinVar CNV (RCV000225330.1) spanning exactly this region (apparently, the tested variant), in one individual, provided by a single submitter was classified as benign (single star) and associated with premature ovarian failure (see screenshot from xxx in which mentioned Chr22 CNV is shown as the only long green line). On the other hand, there is sufficient evidence (175 Pathogenic CNVs fully contained in contrast to 1 Benign CNV fully overlapping this region) to consider this CNV pathogenic, thus MarCNV and ISV classification was also like that.

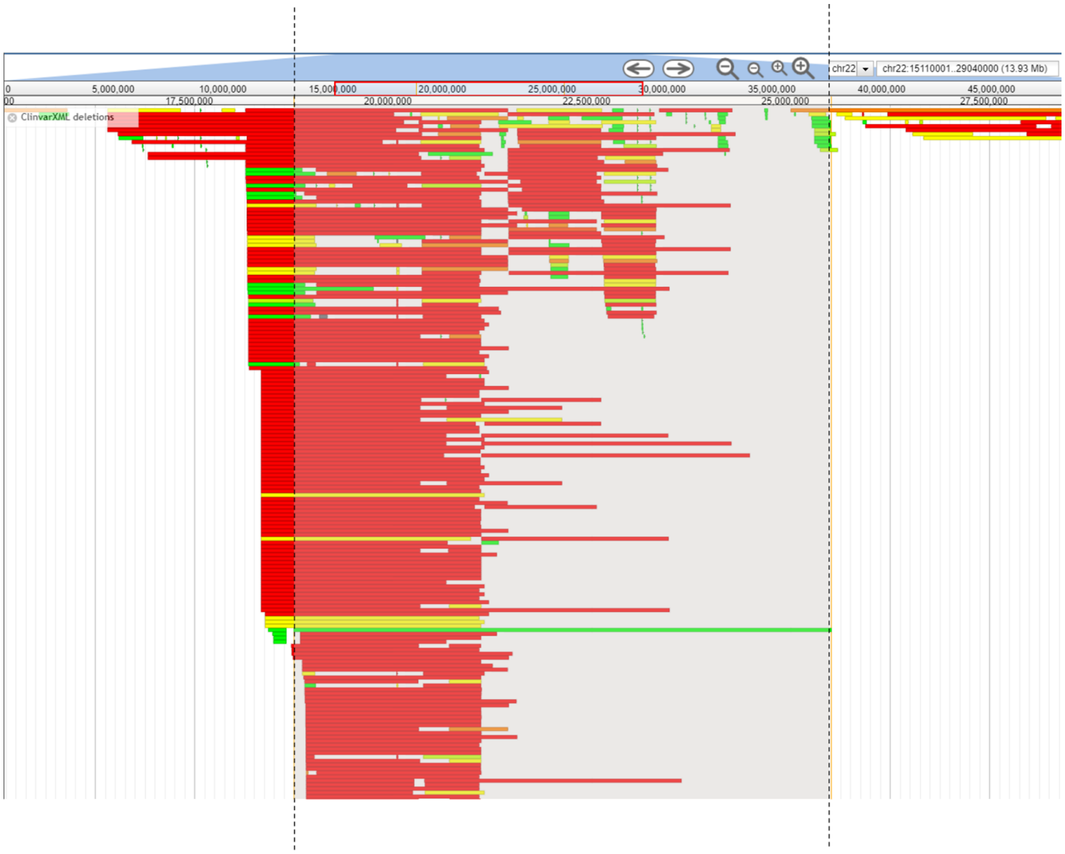


**Losses of chrX:155239112-155691896**; **chrX:77775163-77949550**, and **chr1:16951500-17071209** completely overlap an established haploinsufficient gene/genomic region, so condition 2A. of ACMG criteria was met, and thus MarCNV classified CNVs as Pathogenic. However, most known pathogenic CNVs in these regions are longer and/or overlap only partially, suggesting that the minimal critical region causing pathology is not included in mentioned CNVs coordinates. It is clear that manual correction of automated classification should help in such cases.

**Loss of chr1:145138148-146401981** (dbVar: nsv497843) and **gain of chr2:7495123-87705899** (dbVar: nsv994986) classified in ClinVar as Benign with no assertion criteria provided have been obsoleted and are no longer valid in dbVar database.
